# “Cheater” particles render the VEGAS platform unsuitable for mammalian directed evolution

**DOI:** 10.1101/2022.06.07.495067

**Authors:** Christopher E. Denes, Alexander J. Cole, Minh Thuan Nguyen Tran, Mohd Khairul Nizam Mohd Khalid, Alex W. Hewitt, Daniel Hesselson, G. Gregory Neely

**Author notes:** These authors contributed equally (Christopher E. Denes, Alexander J. Cole). These authors contributed equally (Daniel Hesselson, G. Gregory Neely). Correspondence (D.H.), (G.G.N.).

## Abstract

Directed evolution uses cycles of gene diversification and selection to generate proteins with novel properties. While traditionally directed evolution is performed in prokaryotic systems, recently a mammalian directed evolution system (viral evolution of genetically actuating sequences, or “VEGAS”) has been described. Here we report that the VEGAS system has major limitations precluding its use for directed evolution. The primary technical issue with the VEGAS system is an immediate contamination with “cheater” particles that bypass directed evolution circuits. By sequencing we find these cheater particles contain Sindbis structural genes instead of the intended directed evolution target transgene. These cheaters outcompete the VEGAS transgenes within 2 rounds of transduction but cannot themselves activate synthetic circuits that drive expression of Sindbis structural genes, preventing directed evolution campaigns. Similar results have been obtained in independent labs. Taken together, the VEGAS system does not work as described and, without significant redesign to suppress cheaters, cannot be used for mammalian directed evolution campaigns.

## INTRODUCTION

A new generation of mammalian directed evolution systems have been reported, which exploit viral life cycles to couple mutagenesis of a virally packaged transgene-of-interest with selection for higher fitness variants (Berman et al. 2018; English et al. 2019). In these systems, essential structural components are removed from the viral genome and placed under the control of synthetic gene expression circuits within the host cell and regulated by the transgene-of-interest. Selection operates on natural mutations arising in the transgene during error-prone viral replication that then impact the expression of the transgene-regulated viral structural genes. Each host cell should be infected with a single virus particle each round (i.e. MOI ≤1) so that only the mutation(s) conferring a selective advantage are enriched. Variants that more efficiently drive the synthetic circuit are then overrepresented in the packaged viral particles used for further rounds of evolution (Berman et al. 2018; English et al. 2019). The English, et al. (2019) viral evolution of genetically actuating sequences (VEGAS) system offered a simplified platform for simultaneous mutagenesis and selection using Sindbis virus (SINV). The VEGAS system was designed to link the activity of a transgene to expression of the Sindbis Structural Genome (SSG; comprising Capsid, E3, E2, 6K and E1 genes) (English et al. 2019). For the VEGAS system to support directed evolution, a SINV particle must contain the target transgene, and after transduction should drive expression of the SSG in the next round of evolution.

Here, we find that the VEGAS system does not support viral propagation when performed as described. Similar to English et al. and others (Shapiro et al. 2010; Fayzulin et al. 2005), we found Sindbis virus can efficiently package a transgene and transduce cells, however in the context of the VEGAS system, these particles do not then support the multiple rounds of propagation required for directed evolution. Mechanistically, particles generated using the VEGAS system are immediately contaminated with “cheaters” that contain Sindbis structural genes. These cheaters flood the system, dilute the target transgene, and do not allow directed evolution campaigns.

## RESULTS AND DISCUSSION

In our efforts to implement the VEGAS system, we first attempted to reproduce SSG-dependent replication of SINV (Figure 1B from (English et al. 2019)). To generate Sindbis particles we electroporated BHK-21 cultures with pSinHelper, pSinCapsid and pTSin-EGFP seed mRNA (**Figure S1A)~**. After 24 h, we detected ~10^9^ genome copies (gc)/mL in the supernatant (Round (R)0; **Figure 1A**) and observed bright EGFP fluorescence in most packaging cells (R0; **Figure 1B**). English et al. initiated directed evolution campaigns by transducing target BHK-21 cells with R0 virus at a calculated multiplicity of infection (MOI) of 1 gc/cell (English et al. 2019). Thus, we next transduced target BHK-21 cells in the context of constitutively expressed Sindbis structural genome (CMV-SSG plasmid; +SSG) or control DNA at a calculated MOI of 1 (R1; **Figure 1A**). Importantly, R1 SINV particles were inefficiently generated (<10^6^, >1000x lower titer than R0) and titers were equivalent in samples that received SSG or control DNA (-SSG; 7.35 x 10^6^ ± 2.30 x 10^5^ gc/mL; +SSG, 7.33 ± 4.77 x 10^5^ gc/mL; **Figure 1A**), indicating this system has no dependency on SSG. Moreover, based on GFP expression, SINV particles did not efficiently transduce target cells (**Figure 1B**), and both SSG dependency and transgene delivery are essential requirements for VEGAS directed evolution campaigns.

**Figure 1.**
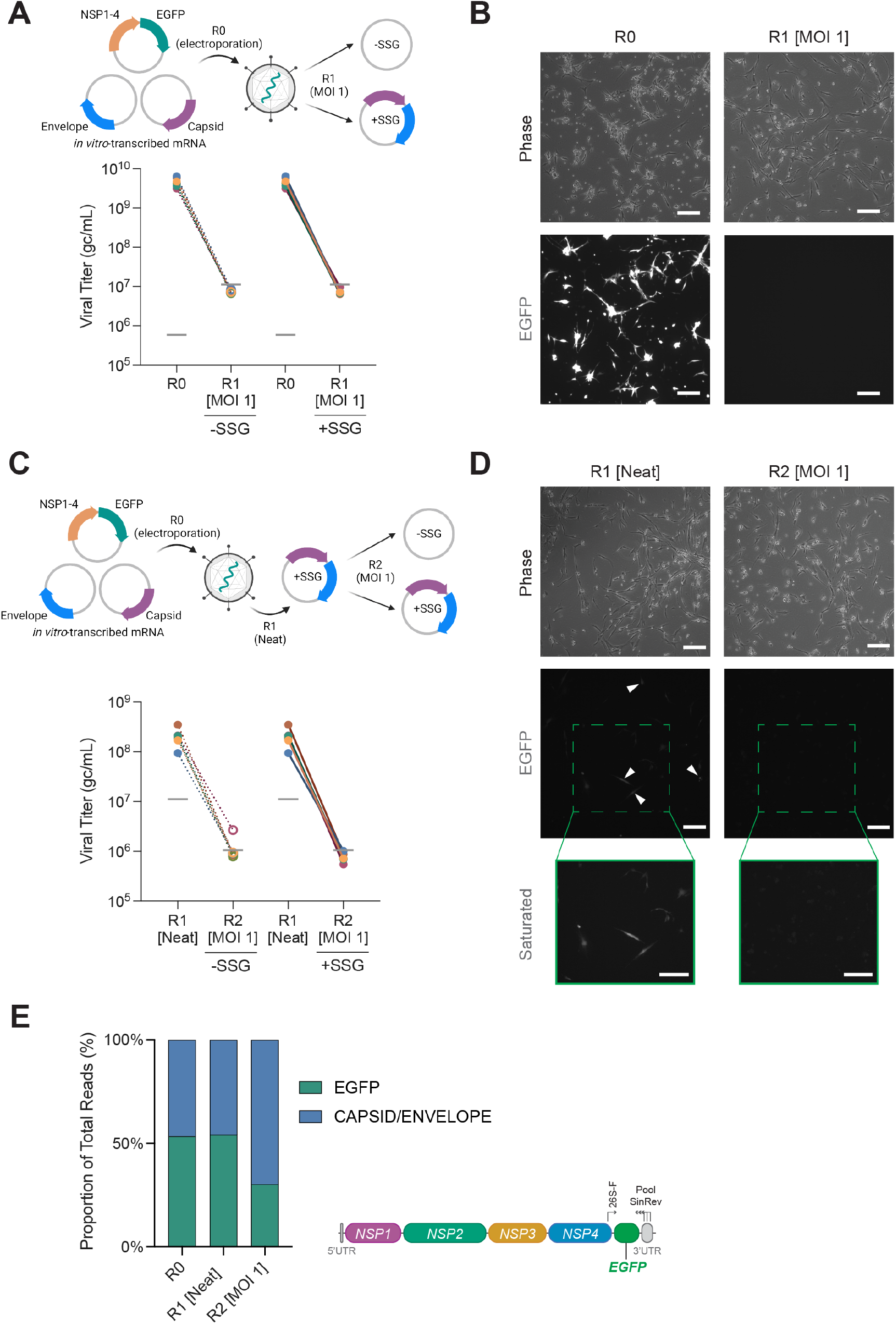
VEGAS generates unproductive transductions. A. Viral titers after packaging and transduction of pTSin-EGFP into control or SSG-expressing BHK-21 cells (N = 6). Horizontal gray bars indicate the batch-specific thresholds for viral detection. B. Phase contrast and EGFP fluorescence images of BHK-21 cells used for packaging (R0) and transduction (R1 [MOI 1]). Scale bars represent 200 μm. C. Viral titers after transduction with undiluted R0 virus into SSG-expressing BHK-21 cells. At R2, viruses were transduced at MOI 1 into control or SSG-expressing BHK-21 cells (N = 6). D. Phase contrast and EGFP fluorescence images of BHK-21 cells transduced with undiluted R0 virus (R1 [Neat]) and MOI 1 at R2. Scale bars represent 200 μm. White arrowheads indicate EGFP-positive cells. Green boxes denote a magnified section of the image with enhanced brightness and contrast to highlight GFP-positive cells. E. Nanopore sequencing of viral RNA isolated from pooled samples in Panels C and D (N = 6). Reads (~100,000 per sample) were aligned to viral reference sequences.

One possible cause for the low viral titers observed at R1 is that transfected host cells failed to express the Sindbis structural genes provided via the CMV-SSG plasmid. Thus, we next examined host cell expression of CMV-SSG at R1. To avoid the possibility of transfected plasmids contributing to high background signal during qPCR, we developed a barcoding method to discriminate between plasmid DNA (pDNA) and expressed mRNA (**Figure S2A**). Using this approach, we confirmed that the CMV-SSG plasmid was highly expressed (>30% of GAPDH, **Figure S2B,C**).

We next addressed possible explanations for the low GFP expression when transduced at an MOI of 1 (**Figure 1B**). We suspected that residual seed mRNA in the supernatant following R0 electroporation could contaminate the R0 output and inflate titers. To test this, we degraded unpackaged seed mRNA by treating R0 and R1 supernatants with RNase A prior to titration. Indeed, RNase A treatment of R0 samples decreased the measured virus titer by 93%, however R1 samples were much less RNase-sensitive (**Figure S3**). This differential sensitivity indicated that residual unpackaged seed RNA was present in the output from R0 confounding accurate MOI calculations.

To determine if inaccurate MOI calculations can explain why the VEGAS system does not support viral propagation after R0, we added undiluted (neat) R0 virus (SINV-EGFP) to CMV-SSG-expressing cells. Transduction with neat virus resulted in 5-10% GFP-positive cells at R1 (arrowheads, **Figure 1D**), indicating that Sindbis packaging and delivery of the target transgene was successful. We then transduced R2 target cells with SINV-EGFP particles at an MOI of 1, however, at R2 we did not observe GFP-positive cells (**Figure 1D**), and output from this transduction was low and not dependent on SSG (**Figure 1C**). These data suggest that bypassing unreliable R1 MOI calculations with neat transductions (and then controlling MOI beyond R1) was still not sufficient to support viral propagation at subsequent rounds of transduction. Overall, the VEGAS system does not support SINV propagation at an MOI of 1, precluding its use for directed evolution campaigns.

To investigate the integrity of the VEGAS system over successive rounds of transduction we performed long-read nanopore sequencing of viral RNA (**Figure 1E**). Using the approach for transgene recovery as described (English et al. 2019), all reads should map to the transgene subjected to directed evolution. Surprisingly, we found only 53.4% of reads mapped to the EGFP transgene in the initial packaged virus (R0; **Figure 1E**) and transgene inclusion was reduced to 30.3% by R2. The remaining reads were from cheater particles that contain envelope and capsid sequences derived from seed RNA provided at R0. We note that the *in vitro-* transcribed VEGAS seed mRNAs (pSinHelper and pSinCapsid) retain components of the SINV NSP1 packaging signal and replication sequences (Weiss, Geigenmüller-Gnirke, and Schlesinger 1994) (**Figure S4**), which would allow seed mRNA to compete with the SINV-EGFP transgene for packaging. These data show that the VEGAS system preferentially packages SINV structural elements, and these cheater particles then outcompete VEGAS transgenes and block directed evolution efforts.

Previous work has shown Sindbis virus can retain a neutral GFP transgene for at least six viral passages at low MOIs (0.1 PFU/cell) (Thomas et al. 2003), however it is possible that in the context of a split VEGAS system, selective pressure for a transgene is required to maintain it across rounds of propagation. To test this possibility, we built a simple synthetic circuit using the DNA-binding domain of serum response factor (SRF, fused to VP64) to drive transgene expression under the control of the serum response element (SRE). We first validated this circuit using SRF-NLS-VP64/SRE-dependent expression of a firefly luciferase reporter (SRE_LUC), which drove luciferase (**Figure 2A)** at levels comparable to (English et al. 2019) (see English et al, **Figure S3B**). Next, we subcloned SRF-NLS-VP64 into the VEGAS pTSin vector and packaged it in a 1:1 ratio with pTSin-EGFP to provide an evolutionarily neutral competitor. Under selection, the EGFP transgene should drop out while the higher fitness SRF-NLS-VP64-containing viruses, capable of driving expression of the Sindbis structural genome, should rapidly dominate the culture. As predicted, the proportion of GFP-positive cells decreased with serial passage (**Figure 2B),** however, this did not translate into efficient SRF-NLS-VP64 viral production at R2 (**Figure 2C**).

**Figure 2:**
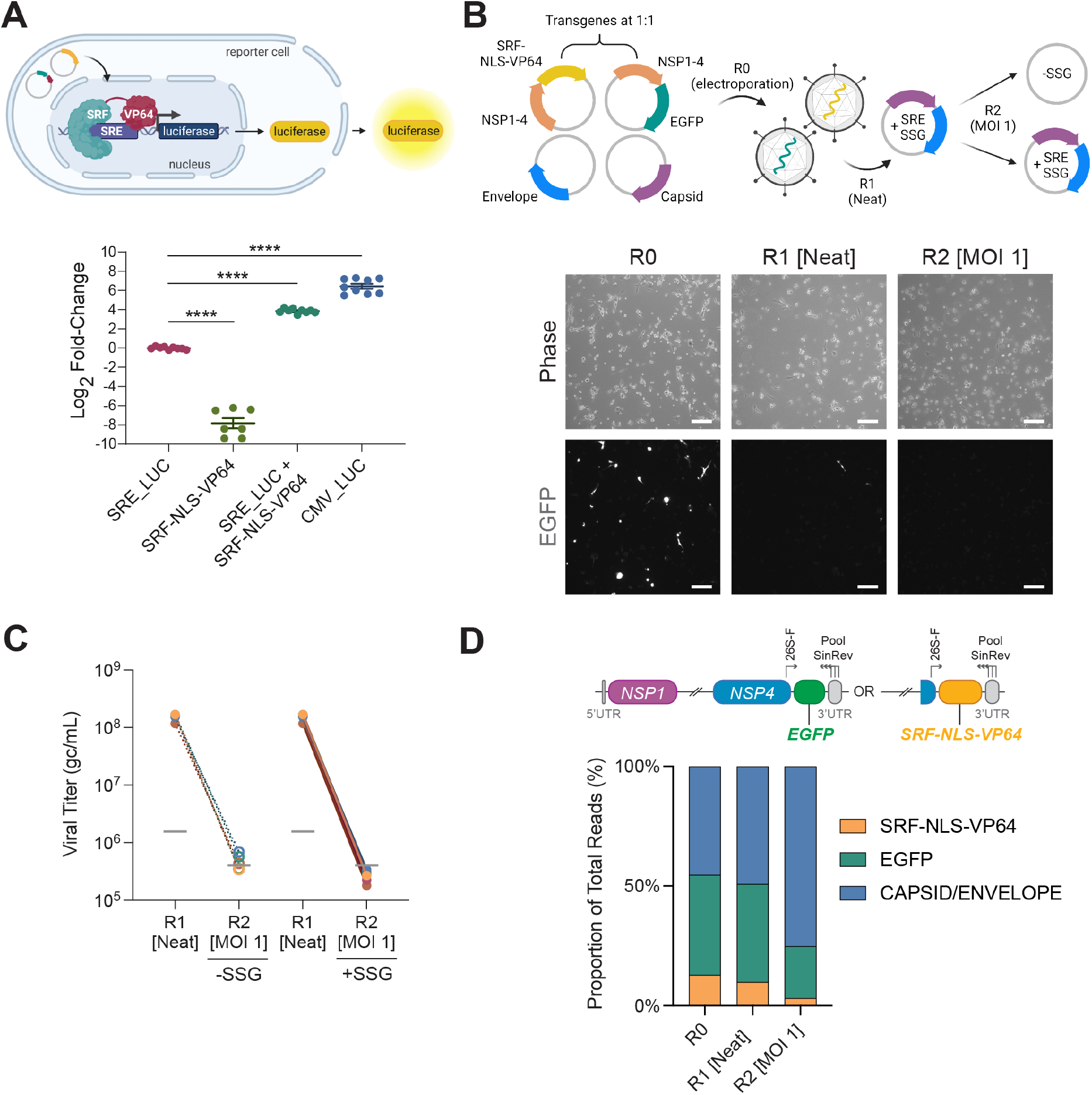
Selective pressure does not rescue system integrity. A. Activation of an SRE-regulated luciferase reporter (SRE_LUC) by SRF-NLS-VP64. Error bars represent mean ± SEM (N = 3, with 3 technical replicates). A Brown-Forsythe and Welch ANOVA test was used to determine statistical significance. <*** *p* < 0.0001. B. Phase contrast and EGFP fluorescence images of BHK-21 cells at R0-R2 for an SRF-NLS-VP64/SRE_SSG circuit. Scale bars represent 200 μm. C. Viral titers of an SRF-NLS-VP64 transgene-carrying virus packaged in a 1:1 ratio with an EGFP-carrying virus (N = 6). D. Nanopore sequencing of viral RNA isolated from pooled samples (N = 6). Reads (~100,000 per sample) were aligned to viral reference sequences.

Importantly, the circuit-inducing SRF-NLS-VP64 transgene did not outcompete the control EGFP transgene (3.2% SRF-NLS-VP64 vs 21.7% GFP by R2), and both transgenes were outcompeted by cheater sequences containing Sindbis structural elements (75.1%, **Figure 2D**). Together, these data show that even under selective pressure, the VEGAS system does not support productive rounds of replication required for directed evolution campaigns. Instead, VEGAS campaigns are flooded with cheater particles containing Sindbis structural genes that prevent the circuit-driven selection required for directed evolution to occur.

While it is possible that undocumented methodological subtleties could explain why VEGAS Sindbis particles fail to propagate across rounds of transduction, we have received feedback from English et al. on the results described here and have thus far not identified any. To generate the Sindbis particles required to initiate VEGAS campaigns, English *et al.* used the NEON electroporator, whereas we used the Amaxa electroporator which also delivers RNA to the cytosol (Ekstrom and Dean 2011) and efficiently generates functional Sindbis particles (Shapiro et al. 2010). Importantly, using a different electroporator does not explain our inability to run VEGAS directed evolution campaigns, since both machines produced Sindbis particles that transduced GFP to R1 target cells with equivalent efficiency (~5-10%). Instead, our data suggests that the VEGAS system is non-functional due to circuit-independent cheater particles that dominate the system.

Our experiments involved VEGAS components generated and made publicly available by English *et al.* when provided (Addgene plasmids #127692–127695). However, we could not obtain the CMV-SSG plasmid from English *et al.* and generated this component ourselves, which we validated and used in **Figure 1**. Moreover, English *et al.* similarly did not provide the VEGAS circuits used in their original work, and thus we generated our own serum response factor circuit used in **Figure 2**. Again, while we confirmed this circuit is functional and could drive expression from the SRE comparable to the circuit used by English *et al.,* we found this circuit could not support viral propagation past R2. A similar inability to use the VEGAS system has been observed independently across our separate laboratories over the three years since the platform was first described.

Importantly, Sindbis virus is pathogenic to humans and has the potential to cause meningitis (Meno et al. 2021; Griffin 2005). Extreme caution should be used when trying to implement the published VEGAS system in a BSL2 environment since a single recombination event between the transgenic SINV genome and the host cell SSG expression plasmid (which share homologous sequences) could produce a replication-competent virus. In our hands we tested for and did not detect replication-competent virus.

While we believe there is a future for Sindbis-based mammalian directed evolution systems, this will require substantial efforts to address critical technical challenges that limit VEGAS utility and, optimally, any repairs or upgrades should be confirmed by independent groups. In summary, although we confirm that packaging and transduction of SINV particles efficiently delivers transgenes to fresh host cells as previously documented (English et al. 2019; Fayzulin et al. 2005; Shapiro et al. 2010), we conclude that due to the prevalence of cheater particles that carry Sindbis structural components instead of target transgenes, the VEGAS system is not suitable for use as a mammalian directed evolution platform.

## METHODS

### Cell Culture

BHK-21 [C-13] cells were purchased from the American Type Culture Collection (#CCL-10). Cells were grown in a humidified 37°C (5% CO2) atmosphere in MEM α (ThermoFisher, #32571101) supplemented with 5% HyClone fetal bovine serum (FBS) (Cytiva Life Sciences, #SH30084.03, AU origin) and 10% tryptose phosphate broth (TPB) (ThermoFisher, #CM0283B), referred to as BHK-21 Growth Medium. During transduction and recovery, cells were maintained in serum-free MEM α with 10% TPB, referred to as BHK-21 Recovery Medium (Serum-Free).

### Molecular Biology and Plasmid Construction

All plasmids were designed in SnapGene® (version 5.3.2). Plasmids were generated by PCR amplification of sequences of interest with Velocity DNA Polymerase (Bioline, #BIO-21099) using primers synthesized by IDT or restriction enzyme digestion with NEB High-Fidelity enzymes. Assembly of amplicons was performed using the NEBuilder HiFi DNA Assembly Master Mix (NEB, #E2621). Assembled products were transformed into NEB® 10-beta Competent *E. coli* (NEB, #C3019) and selected on LB agar plates (ThermoFisher, #22700025) supplemented with 100 μg/ml ampicillin (Sigma-Aldrich, #A9518). Individual colonies grown overnight in liquid LB broth (ThermoFisher, #12795-084) supplemented with 100 μg/ml ampicillin were processed with either the ISOLATE II Plasmid Mini Kit (Bioline, #BIO-52057) for sequence verification, or the PureYield™ Plasmid Maxiprep System (Promega, #A2393) for transfection and *in vitro* transcription applications. Plasmid constructs were verified by restriction digestion and Sanger sequencing at the Australian Genome Research Facility (AGRF). A list of plasmids used and generated in this study is available in Supplementary File 1 – Plasmid List.

### mRNA synthesis

mRNA for electroporation and transfection was produced using the mMESSAGE mMACHINE^TM^ SP6 Transcription Kit (ThermoFisher, #AM1340) from XbaI-linearized SP6-driven plasmids as outlined previously (English et al. 2019). mRNA concentrations were calculated using the Qubit™ RNA BR Assay Kit (ThermoFisher, #Q10210). mRNA integrity was assessed by gel electrophoresis (**Figure S1**). mRNA was frozen at-80°C immediately after transcription as per (English et al. 2019).

### Packaging of SINV Particles (referred to as Round 0 (R0))

1 x 10^6^ BHK-21 cells were electroporated with a total of 7.8 μg of mRNA (1:1:1 of pSinHelper, pSinCapsid and pTSin-EGFP/pTSin-SRF-NLS-VP64) using Amaxa 2B (Lonza) as per the manufacturer’s instructions for BHK-21 cells. The Amaxa electroporator efficiently delivers SINV RNA into the cell cytosol (Ekstrom and Dean 2011), allowing for the packaging of mature SINV particles (Shapiro et al. 2010). Electroporated cells were plated in BHK-21 Recovery Medium (Serum-Free). Virus-containing supernatants were collected 24 hours after mRNA delivery and centrifuged at 500 *g* for 5 minutes to pellet cellular debris. Clarified supernatants were collected for titration and subsequent transduction experiments.

### Viral Titration

Viral supernatants were titrated as outlined by English and colleagues (English et al. 2019). Following collection and clarification, undiluted SINV-containing supernatants were combined with the TaqMan™ Fast Virus 1-Step Master Mix (ThermoFisher, #4444434) and the appropriate primer-probe sets in technical triplicates. Plates were run on a QuantStudio™ 7 Flex Real-Time PCR System (ThermoFisher). As described previously, serially diluted *in vitro*-transcribed pTSin mRNA was used as the standard curve (ranging between 10^3^-10^7^ genome copies (gc) per mL) (English et al. 2019). Standard curves were used to determine viral titers in gc/mL. Primer sequences are listed in Supplementary File 2 – Primer List.

### Transduction of Viral Particles (Round 1 (R1)-onwards)

BHK-21 cells were seeded in 6-well plates at 6.5 x 10^4^ cells/well, incubated for 24 hours, and transfected with a total of 2.5 μg of SSG-encoding DNA/well using *TransIT-2020* Transfection Reagent (Mirus Bio, #MIR5400) following the manufacturer’s recommendations. 6 hours post-transfection, cells were rinsed twice with DPBS before viral inoculum (either undiluted or diluted to a calculated MOI of 1 gc/cell) was applied in a 380 μl volume of BHK-21 Recovery Medium (Serum-Free). Cells were incubated with virus for 1 hour and rinsed twice with DPBS before 1.5 mL BHK-21 Recovery Medium (Serum-Free) was added for a further 23 hours of incubation. Virus-containing supernatants were collected and processed for titration and transduction experiments as described in *Packaging of SINVParticles.*

We note that recombination between the SINV genome and homologous sequences in the SSG 3’UTR (**Figure S4)** could produce competent viral populations. While this has not been observed in our independent labs, we urge responsible research practices and approvals for working in appropriate Biosafety Level 2 facilities, and the application of appropriate testing protocols for determining replication competency. Assaying viral titer amplification on cells ± SSG elements in parallel can be used to quickly ascertain structural element dependency status.

### RNase Digestion

To test if inflated R0 viral titers were a result of residual packaging mRNA, pooled viral supernatants were digested with 0 or 400 ng RNase A (Macherey-Nagel, #740505.50) for 4 hours at 37°C. Samples were then incubated for 20 minutes at room temperature with 80 units of RNaseOUT (ThermoFisher, #10777019). Samples were titrated as described in *Viral Titration.* RNase A treatment was not applied to any samples used for subsequent transduction assays.

### Barcoded qPCR

RNA was extracted using the FavorPrep™ Blood/Cultured Cell Total RNA Mini Kit (Fisher Biotec, #FABRK 001-1) and 400 ng of cDNA was synthesized using the iScript™ Select cDNA Synthesis Kit (Bio-Rad Laboratories, #1708897) on a Mastercycler® nexus PCR Thermal Cycler (Eppendorf): 42°C for 60 min, 85°C for 5 min. cDNA was produced using two primers: a specific primer for hamster GAPDH, and a SINV structural genome primer containing a barcode. To remove residual primers, cDNA was bound to AMPure XP magnetic beads (Beckman Coulter, #A63881) in 0.75 M LiCl, 20% PEG 8000 buffer at a 1:0.9 DNA:buffer volume ratio, and washed twice with 70% ethanol. qPCR was performed using SYBR™ Select Master Mix (ThermoFisher, #4472908) following manufacturer’s recommendations in triplicate on a QuantStudio™ 7 Flex Real-Time PCR System (ThermoFisher). Primer sequences are listed in Supplementary File 2 – Primer List.

### Luciferase Assay

BHK-21 cells were seeded in 96-well plates at 2200 cells/well. Cells were transfected with a total 90 ng of plasmid constructs using *TransIT-2020* Transfection Reagent following the manufacturer’s recommendations. After 6 hours, cells were washed twice with DPBS and switched to BHK-21 Recovery Medium. At 24 hours post-transfection, luciferase activity was assessed using the Dual-Glo Luciferase Assay System (Promega, #E2940) following manufacturer’s recommendations in black-bottom plates. Both Renilla and firefly luminescence were measured on an Infinite M1000 PRO microplate reader (Tecan). Raw firefly luminescence values were normalized to Renilla luminescence and log2-transformed.

### Transgene Isolation

Sextuplicate viral supernatants from each round of viral replication were pooled in equal ratios. Viral RNA from 400 μl aliquots was isolated with the MagMAX™ Viral RNA Isolation Kit (ThermoFisher, #AM1939) as per the manufacturer’s recommendations. Transgene sequences, situated between nsP4 and the viral 3’ untranslated region (UTR) were reverse-transcribed and PCR-amplified using the SuperScript™ IV One-Step RT-PCR System (ThermoFisher, #12594025) and the 26S-F primer and pooled SinRev primer. Primer sequences are listed in Supplementary File 2 – Primer List. Amplicons of the appropriate size range for the transgene of interest were then gel extracted for nanopore sequencing using the ISOLATE II PCR and Gel Kit (Bioline, #BIO-52060) (**Figure S5**).

### Nanopore Sequencing

#### Sample Processing

Isolated transgene DNA was processed to generate libraries for full-length transgene sequencing using Oxford Nanopore Technologies (ONT) Flongle flow cells (ONT, FLO-FLG001). Samples were prepared for sequencing according to the ONT protocol ‘Amplicons by Ligation’ (version ACDE_9064_V109_REVP_14AUG2019). 200 fmol of DNA (determined following quantification with the Qubit™ dsDNA HS Assay Kit) was processed using the NEBNext Companion Module for ONT Ligation Sequencing (NEB, #E7180). Sequencing adapters were added using the Ligation Sequencing Kit (ONT, #SQK-LSK109).

#### Sequencing and Basecalling

Up to 40 fmol of DNA library was loaded onto ONT Flongle flow cells (R9.4.1) in a MinION Mk1B Sequencer fitted with a Flongle Adapter (ONT). Sequencing was performed using MinKNOW (ONT, version 4.2.8) under default parameters and the following specified inputs: kit used, SQK-LSK109; 0.5 hours between MUX scans; basecalling, disabled. A minimum of 120,000 raw reads were obtained for each sample. Raw FAST5 files were processed using Guppy (version 4.5.2) with the minimum q-score set to 7.0. Raw FAST5 and basecalled FASTQ reads were deposited at the European Nucleotide Archive (PRJEB47639).

#### Alignment

Quality-filtered base-called reads (in FASTQ format) were processed using EPI2ME Desktop Agent (ONT, version 3.3.0.1031). Reference files, consisting of complete transgene-coding sequences or helper-and capsid-coding sequences, were uploaded using the Fasta Reference Upload workflow (v2021.07.15). For each sample, 100,000 reads were aligned to reference files using the Fastq Custom Alignment workflow (v2021.03.25) using default parameters.

#### Microscopy

Phase contrast and EGFP fluorescence images were captured at 5X magnification on an Axio Vert.A1 FL(Zeiss) fitted with an AxioCam ICM1 camera (Zeiss 60N-C 2/3” 0.63X adapter). EGFP images were obtained with a BP475/40 excitation and BP530/50 emission filter (FT500 beam splitter). Images were collected using Zen 2 Blue Edition (Zeiss, version 2.0.0.0).

#### Statistics

Statistical analyses were performed in GraphPad Prism 9.2.0 for Mac, GraphPad Software, San Diego, California USA, www.graphpad.com. Fold-changes in luciferase activity were statistically analysed with a Brown-Forsythe and Welch ANOVA test, assuming Gaussian distribution and unequal standard deviations. Means were compared to the control baseline mean. qPCR data was analysed using multiple student t-test. *p-*values < 0.05 were considered significant. All data were plotted as mean ± SEM.

## ACKNOWLEDGMENTS

A.W.H. was supported by National Health and Medical Research Council (NHMRC) Practitioner Fellowship GNT1150144. D.H. is funded by NHMRC grant GNT1185002 and the Centenary Institute. G.G.N. is funded by NHMRC grants GNT1107514, GNT1158164, GNT1158165, GNT1185002, the NSW Ministry of Health, and a kind donation from Dr. John and Anne Chong. Figure schematics were created with BioRender.com.

## AUTHOR CONTRIBUTIONS

Conceptualization, CED, AJC, DH, GGN; Methodology, CED, AJC, MTNT, DH; Investigation, CED, AJC; Resources, AWH, GGN; Data analysis, CED, AJC, MKNMK, MTNT, DH; Writing-original draft, CED, AJC, MTNT, DH; Writing-review & editing, CED, AJC, MTNT, MKNMK, AWH, DH, GGN; Supervision and project administration, AWH, DH, GGN; Funding acquisition, AWH, DH, GGN.

## INCLUSION AND DIVERSITY

One or more of the authors of this paper self-identifies as a member of the LGBTQ+ community.

## SUPPLEMENTAL FIGURE LEGEND

**Figure S1.**
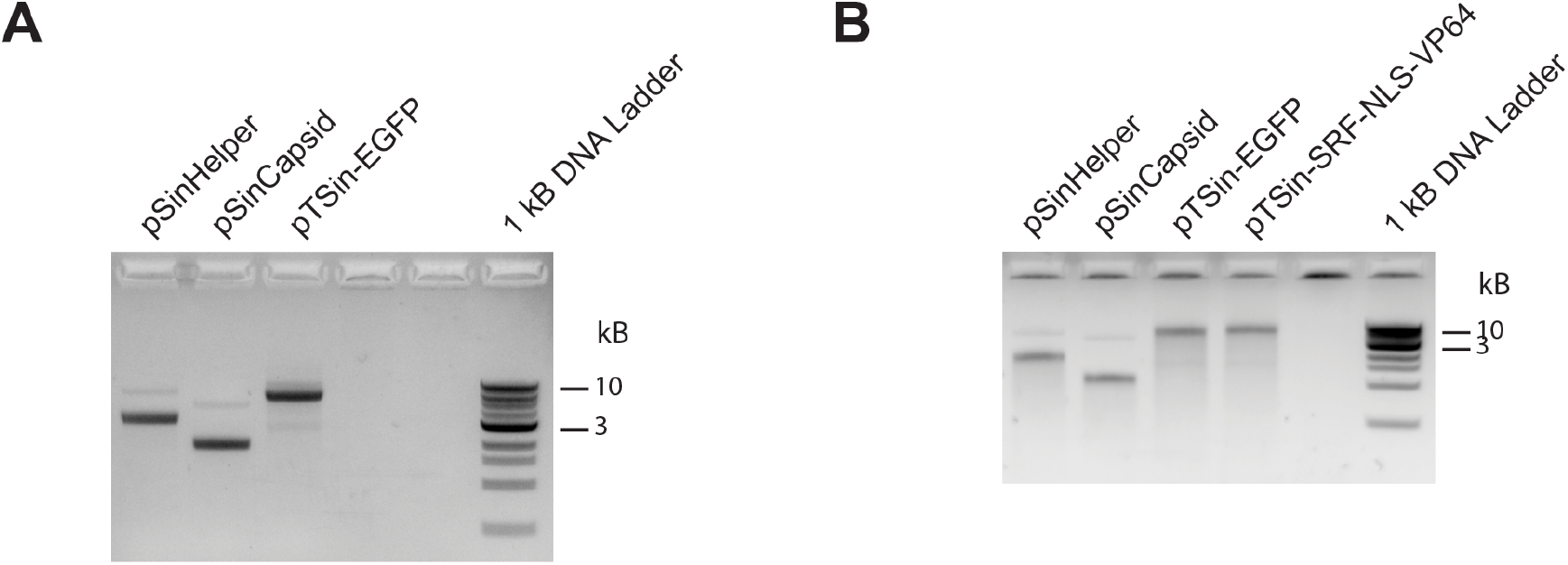
Integrity of *in vitro*-transcribed mRNA. A. Gelelectrophoresis of VEGAS *in vitro*-transcribed mRNA. B. Gelelectrophoresis of *in vitro*-transcribed mRNA used for SRF-NLS-VP64 selection circuit.

**Figure S2.**
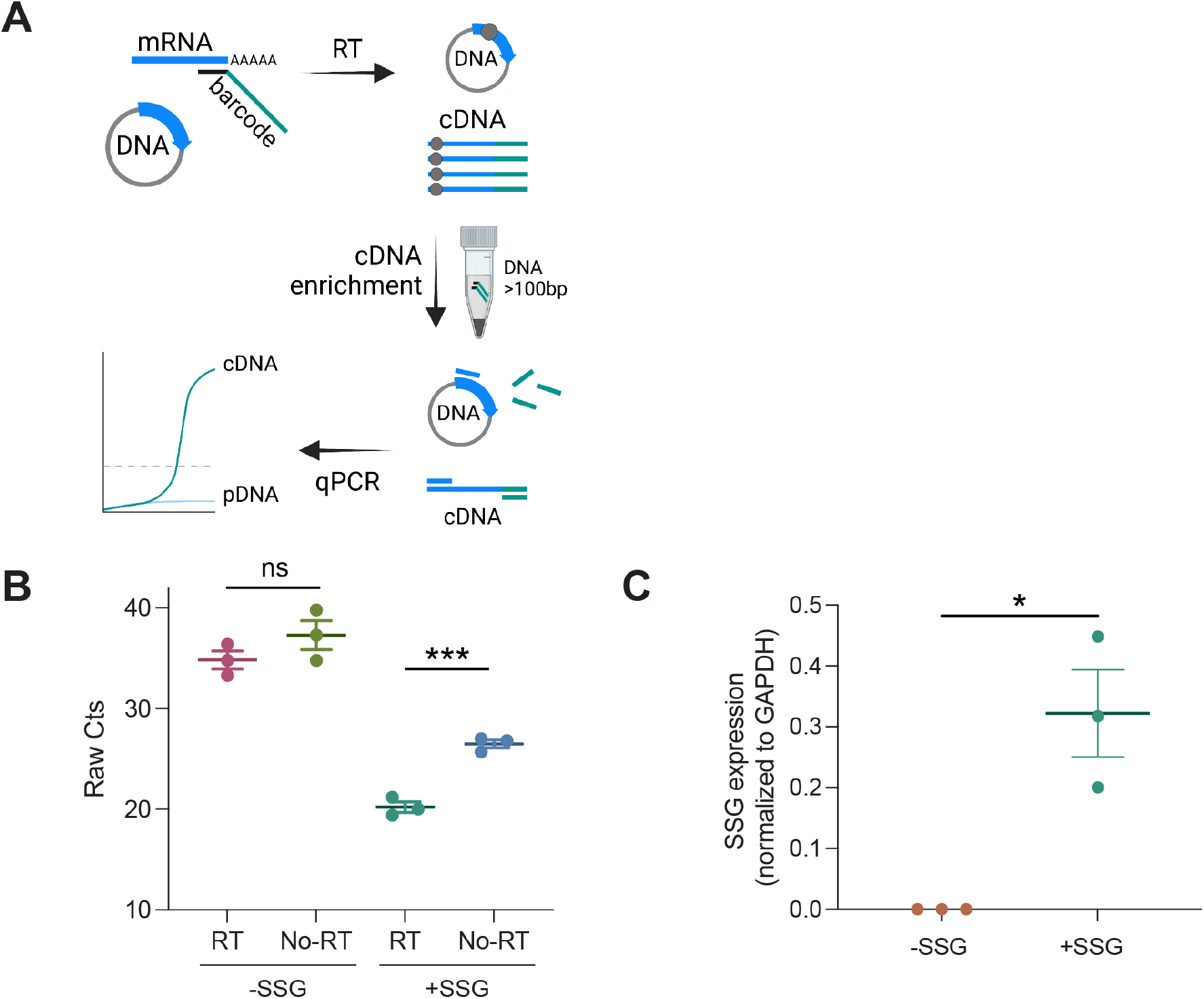
Expression of the SSG in transfected BHK-21 cells. A. Schematic of the barcoded qPCR method used to discriminate mRNA from pDNA during transient transfection. A barcoded SSG-specific primer was used for reverse transcription. Unincorporated barcoded primers were depleted by bead clean-up (indicated by gray circles). qPCR was performed using an SSG-specific forward primer and a barcode-specific reverse primer to discriminate between SSG cDNA and residual pDNA. B. Raw Ct values from barcoded qPCR for SSG expression of the CMV-SSG plasmid in BHK-21 cells, ± reverse transcription (RT). t-tests were used to determine statistical significance. *\*\*\*p* < 0.001, ns = non-significant. C. SSG expression was calculated relative to GAPDH. Error bars represent mean ± SEM (N = 3). t-tests were used to determine statistical significance. \**p* < 0.05.

**Figure S3.**
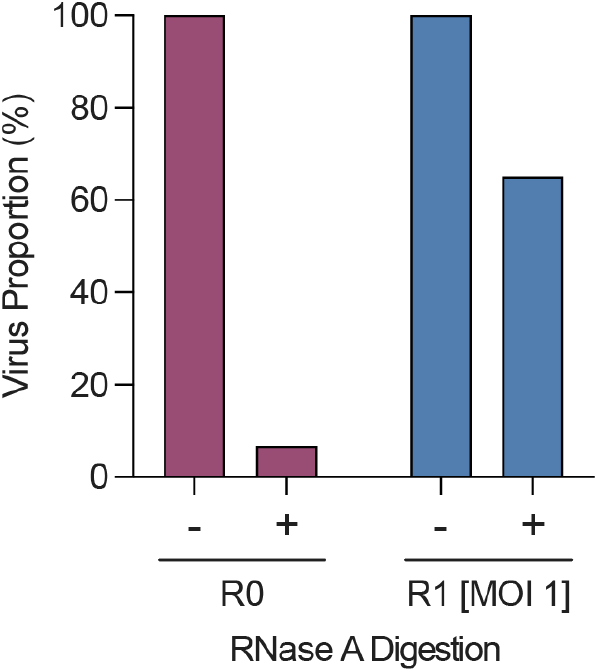
RNase A-digested R0 and R1 viruses. Pooled SINV from R0 and R1 (N = 6, produced on +SSG BHK-21 cells) were treated with RNAse A and titrated.

**Figure S4.**
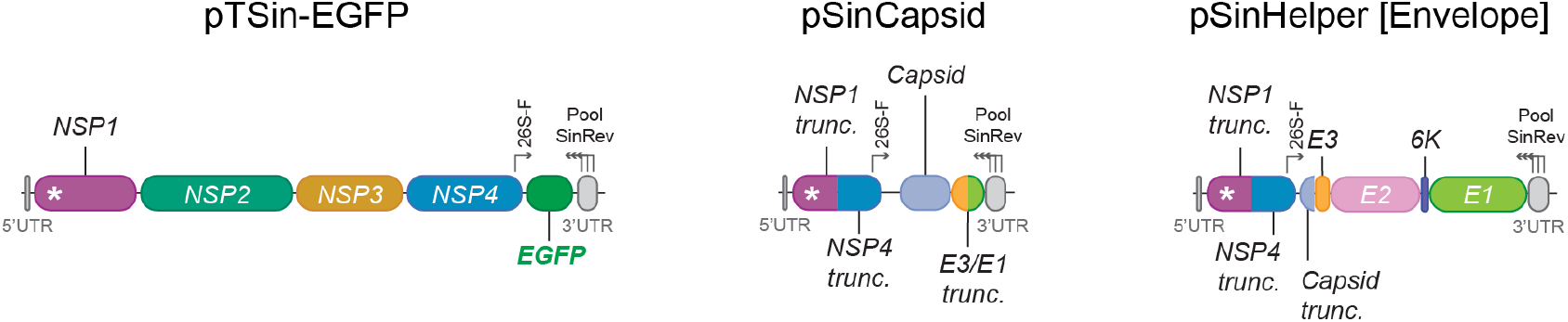
Packaging signal alignment in seed mRNA sequences. Schematic showing the position of the SINV packaging signal within VEGAS seed mRNAs (asterisks). Genes and truncations drawn to scale.

**Figure S5.**
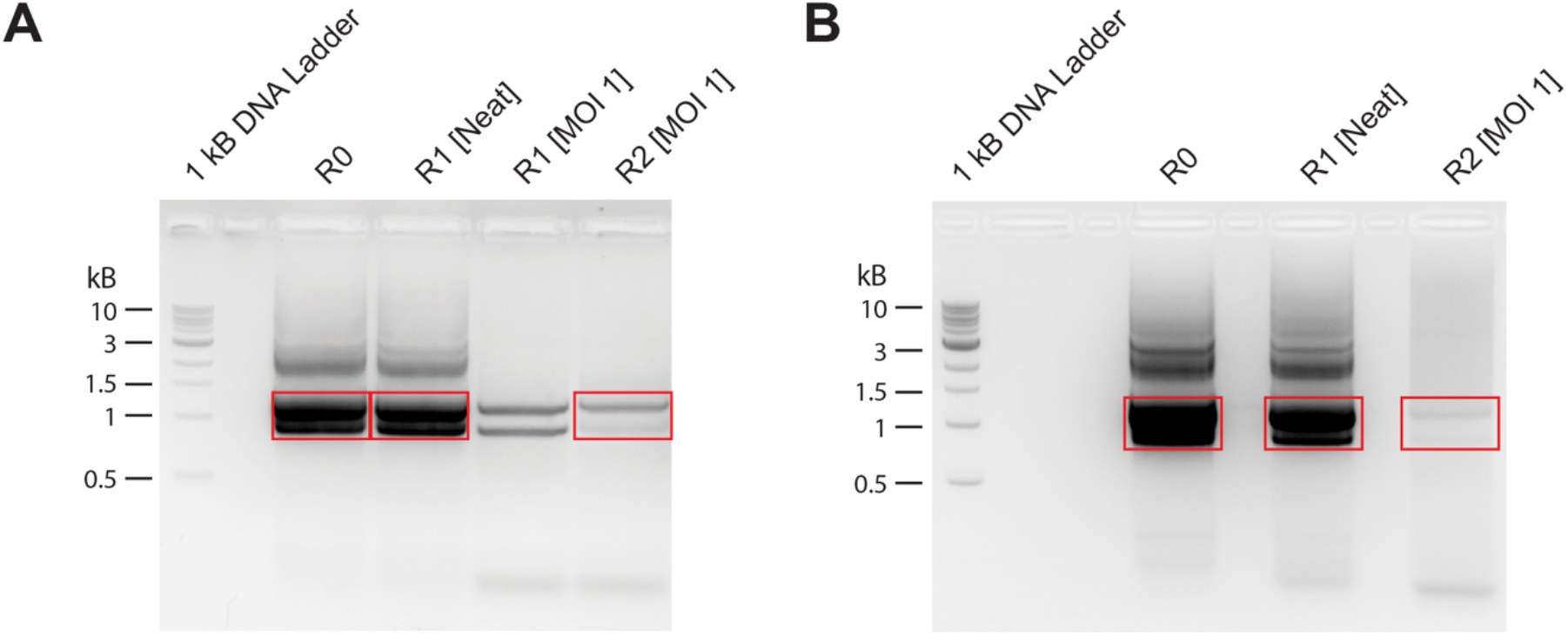
Isolated transgene RT-PCR amplicons for nanopore sequencing. A. Transgenes from electroporated VEGAS viruses (from Figure 1) were amplified by RT-PCR (boxed in red) and gel extracted for nanopore sequencing. B. Transgenes from the SRF-NLS-VP64/SRE_SSG circuit (from Figure 3C) were amplified by RT-PCR (boxed in red) and gel extracted for nanopore sequencing.

## KEY RESOURCES TABLE

| REAGENT or RESOURCE | SOURCE | IDENTIFIER |
| --- | --- | --- |
| **Bacterial Strains** |  |  |
| NEB® 10-beta Competent *E. coli* (High Efficiency) | NEB | C3019 |
| **Chemicals, Peptides, and Recombinant Proteins** |  |  |
| Ampicillin sodium salt | Sigma-Aldrich | A9518 |
| RNase A | Macherey-Nagel | 740505.50 |
| **Critical Commercial Assays** |  |  |
| Dual-Glo® Luciferase Assay Kit | Promega | E2940 |
| FavorPrep™ Blood/Cultured Cell Total RNA Mini Kit | Fisher Biotec | FABRK 001-1 |
| iScript™ Select cDNA Synthesis Kit | Bio-Rad Laboratories | 1708897 |
| ISOLATE II PCR and Gel Kit | Bioline | BIO-52060 |
| ISOLATE II Plasmid Mini Kit | Bioline | BIO-52057 |
| Ligation Sequencing Kit | Oxford Nanopore Technologies | SQK-LSK109 |
| MagMAX™ Viral RNA Isolation Kit | ThermoFisher | AM1939 |
| mMESSAGE mMACHINE™ SP6 Transcription Kit | ThermoFisher | AM1340 |
| NEBNext® Companion Module for ONT Ligation Sequencing | NEB | E7180 |
| NEBuilder® HiFi DNA Assembly Master Mix | NEB | E2621 |
| PureYield™ Plasmid Maxiprep System | Promega | A2393 |

| **REAGENT or RESOURCE** | **SOURCE** | **IDENTIFIER** |
| --- | --- | --- |
| Qubit™ dsDNA HS Assay Kit | ThermoFisher | Q32851 |
| Qubit™ RNA BR Assay Kit | ThermoFisher | Q10210 |
| SuperScript™ IV One-Step RT-PCR System | ThermoFisher | 12594025 |
| SYBR™ Select Master Mix | ThermoFisher | 4472908 |
| TaqMan™ Fast Virus 1-Step Master Mix | ThermoFisher | 4444434 |
| *TransIT*®-2020 Transfection Reagent | Mirus Bio | MIR5400 |
| Velocity DNA Polymerase | Bioline | BIO-21099 |
| **Deposited Data** |  |  |
| European Nucleotide Archive | This paper | PRJEB47639 |
| **Experimental Models: Cell Lines** |  |  |
| Hamster: BHK-21 [C-13] | ATCC | CCL-10 |
| **Software and Algorithms** |  |  |
| EPI2ME Desktop Agent (Version 3.3.0.1031) | Oxford Nanopore Technologies | https://www.nanoporetech.com |
| GraphPad Prism (Version 9.2.0) | GraphPad Software | https://www.graphpad.com/scientific-software/prism/ |
| Guppy (Version 4.5.2) | Oxford Nanopore Technologies | https://www.nanoporetech.com |
| MinKNOW (Version 4.2.8) | Oxford Nanopore Technologies | https://www.nanoporetech.com |
| SnapGene® (Version 5.3.2) | Insightful Science | https://www.snapgene.com |
| UMIC-seq | GitHub | https://github.com/fhlab/UMIC-seq |
| Zen 2 Blue Edition (Version 2.0.0.0) | Zeiss | https://www.zeiss.com |
| **Recombinant DNA** |  |  |
| See Supplementary File 1 - Plasmid List. Plasmid maps are available upon request. | N/A | N/A |
| **Oligonucleotides** |  |  |
| See Supplementary File 2 - Primer List. | N/A | N/A |
| **Other** |  |  |
| 0.45 µm filter | Merck Millipore | SLHV033RS |
| 6-well plate | Corning | 3516 |
| 96-well plate | Corning | 3596 |
| 96-well plate, black-bottom | Interpath | 655209 |
| AMPure XP Magnetic Beads | Beckman Coulter | A63881 |
| DMEM, high glucose | ThermoFisher | 11965118 |
| DPBS | Sigma-Aldrich | D8537 |
| Fetal Bovine Serum (HyClone), AU origin | Cytiva Life Sciences | SH30084.03 |
| Flongle Flow Cell (R9.4.1) | Oxford Nanopore Technologies | FLO-FLG001 |
| LB Agar | ThermoFisher | 22700025 |
| LB Broth | ThermoFisher | 12795-084 |
| MEM $\alpha$ with nucleosides | ThermoFisher | 32571101 |
| Opti-MEM Reduced Serum Medium<br>(GlutaMAX Supplement) | ThermoFisher | 51985034 |
| RNaseOUT Recombinant Ribonuclease<br>Inhibitor | ThermoFisher | 10777019 |
| Tryptose Phosphate Broth | ThermoFisher | CM0283B |
| XbaI | NEB | R0145 |

